# Selective Effects of Ongoing Alpha-Band Activity on Magno- and Parvo-Mediated Detection

**DOI:** 10.1101/2024.07.28.605527

**Authors:** April Pilipenko, Jason Samaha

## Abstract

Spontaneous fluctuations in cortical excitability, as reflected in variation in occipital alpha-band activity (8-12 Hz), have been shown to explain trial-to-trial variability in perception. Specifically, observers typically report seeing a stimulus more often during states of weak alpha power, likely due to a shift in detection criterion. However, prior work has paid little attention to the specific stimulus properties mediating detection. In early vision, different stimulus properties are preferentially processed along the magnocellular (MC) and parvocellular (PC) pathways, which vary in their preference for spatial and temporal frequency content and chromatic information. The goal of this study was to understand how spontaneous alpha power affects the detection of stimuli which are preferentially processed by either the MC or PC pathway. To achieve this, we used the “Steady/Pulsed Paradigm’’ which presented a brief, near-threshold stimulus in two conditions intended to bias processing to one or the other pathway. Our results showed an interaction effect of pre-stimulus alpha power on detection between the two conditions. While weak alpha power was predictive of seeing the stimulus in the steady condition (MC-biased), no significant effect was found in the pulsed condition (PC-biased). This interaction was driven by a selective alpha-related criterion shift in the steady condition, with no effect of pre-stimulus alpha on sensitivity (d’) in either condition. Our results imply that alpha oscillations may differentially regulate excitability in the MC and PC pathways.

## Introduction

Alpha-band oscillations (8-12 Hz) are one of the most prominent features of the awake electroencephalography (EEG) signal in humans. Alpha amplitude fluctuations at posterior cortical sites have repeatedly been associated with visual detection, such that observers are more likely to detect a near-threshold visual stimulus during a period of weak alpha power (Ergenoglu et al., 2004; Mathewson et al., 2009; Busch et al., 2009; for a review: Iemi et al., 2017). More recent research from the past decade has used signal detection theory (SDT) in order to disentangle detection changes that arise from either sensitivity (d’) or bias (criterion or c) by incorporating stimulus-absent trials. A change in sensitivity is characterized by better (or worse) discrimination between states of stimulus-presence and stimulus-absence whereas a change in criterion implies an overall bias to respond a particular way (either “seen” or “unseen”). A number of studies looking at the effect of alpha using the SDT framework suggest a shift in criterion between states of strong and weak alpha power (Lange et al., 2013; Limbach & Corballis, 2016; Craddock et al., 2017; Iemi et al., 2018). Specifically, observers tend to become more liberal when alpha is weak and report perceiving targets more frequently regardless of whether the target was present, with no corresponding change in d’. In discrimination tasks, a similar finding emerges whereby participants report higher confidence or visibility during states of weak alpha, typically alongside no changes in accuracy (Samaha et al., 2017; Wöstmann et al., 2019; Benwell et al., 2022; Samaha et al., 2022).

Based on the above findings, alpha power has been hypothesized to reflect states of visuocortical excitability, known as the baseline sensory excitability model (BSEM; Iemi et al. 2017; Samaha et al., 2020). The shift in criterion can be interpreted as a global increase in neuronal excitability whereby neural populations representing signal and noise are both boosted, making signal and noise more likely to surpass an observer’s detection criterion. Accordingly, alpha power has been presumed to affect all incoming visual content indiscriminately; however, there is currently insufficient evidence to fully support this claim. One specific area where we lack understanding is in the effect of alpha on pathway-specific visual information.

A major distinction in early visual processing is between the magnocellular (MC) and parvocellular (PC) pathways (Maunsell et al., 1990; Masri et al., 2020; Solomon, 2021). The MC pathway, which more heavily innervates peripheral vision, responds to fast, transient events and has relatively large receptive fields which are somewhat more selective to lower spatial frequencies (Solomon et al., 2006). The MC pathway is believed to preferentially project into the dorsal (“where”) stream, which plays a crucial role in processing movement and location (Breitmeyer, 2014; Milner & Goodale, 2008). In contrast, the PC pathway, which is concentrated more foveally, processes color, has smaller receptive fields suitable for higher spatial frequencies, responds to sustained visual events (Solomon et al., 2006), and is thought to have greater innervations to the ventral (“what”) pathway, which has classically been associated with object recognition (Breitmeyer, 2014; Milner & Goodale, 2008).

Much prior work in humans has attempted to study the MC and PC pathway psychophysically, by leveraging these differences in feature processing in order to create conditions which bias stimulus processing to either the MC or PC pathway (Edwards et al., 2021). The “Steady and Pulsed Pedestals” (SPP) paradigm is a widely-used protocol developed and extensively tested since the mid-1990s (Pokorny & Smith, 1997; for a review: Pokorny, 2011). The SPP paradigm relies on the differences in response gain properties of the two pathways and has two main conditions, an MC-mediated “steady (pedestals)” condition and a PC-mediated “pulsed (pedestals)” condition. In the pulsed condition there is a gray background of constant luminance and during the stimulus presentation, a set of luminance pedestals are presented alongside the target (see Figure 1A). The sudden onset of the high-contrast pedestals is thought to saturate the response of magnocellular neurons and bias detection towards the PC pathway. The steady condition is nearly identical to the pulsed condition – it consists of the same uniform, gray background and target stimulus – the crucial difference is that the set of luminance pedestals are consistently shown throughout the entire intertrial and stimulus presentation interval (see Figure 1A). The constant luminance difference of the pedestals to the background elicits a sustained response from the PC pathway, which is thought to lead to adaptation in the PC pathway, thus biasing detection to the MC pathway. Decades of use of this paradigm have shown that it captures key tuning properties of the two pathways (Pokorny, 2011) and one advantage of the SPP paradigm over other psychophysical approaches to pathway separation is that target stimulus properties can be held constant across tasks while only varying the adaptation history.

**Figure 1:**
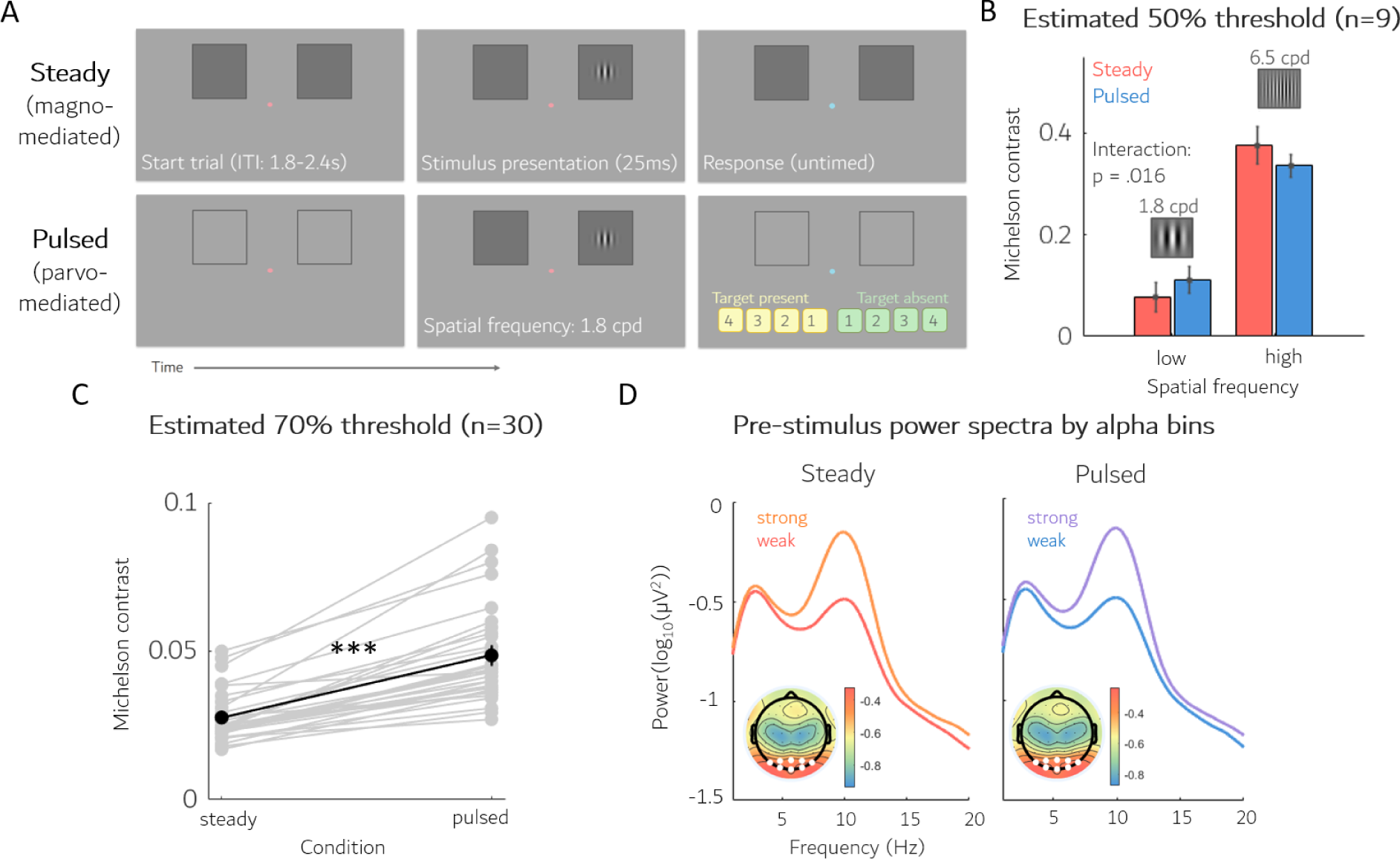
**A**) A schematic of the experimental task. This design implements a variation of the “Steady and Pulsed Pedestals” paradigm (Pokorny & Smith, 1997) wherein participants had to detect a target Gabor patch that occurred atop a steady luminance pedestal (thought to adapt the PC pathway) or atop with a pulsed luminance pedestal (thought to saturate the MC pathway). **B**) Pilot data demonstrates that changing the spatial frequency of the Gabor target modulates detection thresholds in a way that is congruent to the feature preference of a given pathway. Namely, when the spatial frequency is relatively low, a lower threshold is needed for the MC-mediated steady condition. This effect flips when using a high spatial frequency Gabor, requiring a lower threshold for the PC-mediated pulsed condition. **C**) The detection threshold values of the main study. As predicted, there is a significant increase in the threshold level for the pulsed condition compared to steady, given the target Gabor spatial frequency of 1.8. **D**) The average power spectra for weak and strong alpha trials in steady and pulsed conditions. Alpha power was assigned to bins using a median split for both conditions. The grand average topoplot for 8-12 Hz activity. The 8 white circles represent the electrodes with the strongest alpha-band power and were averaged over when computing single-trial alpha power (see *Methods*). Error bars are +/- 1 SEM.

The current study is interested in how the effects of alpha power on detection might manifest differently in the MC and PC pathways. A large motivator for this was the observation that many studies looking at alpha on perception deploy stimuli whose features favor MC-detection. For example, stimuli are often short with transient onsets and offsets (Mathewson et al., 2009; Chaumon & Busch, 2014), achromatic and vary in either luminance or contrast (Limbach & Corballis, 2016; Benwell et al., 2017), and have low spatial frequencies (Iemi & Busch, 2018; Balestrieri & Busch, 2022). To date, no study has investigated whether these common stimulus properties may be a mediating factor in the results. Using the SPP paradigm, we collected EEG recordings and simultaneous detection and confidence judgments from 30 participants. We extracted single-trial alpha power estimates in order to assess the relationship that alpha has on detection between the MC and PC pathway. We found an interaction effect of alpha on detection depending on the condition, suggesting that alpha may not be broadly affecting all visual content, and instead may have a selective effect on perception mediated by the MC pathway.

## Materials and methods

### Subjects

We recruited 33 participants with normal or corrected vision from the University of California, Santa Cruz research participant pool. Participants were awarded 3.5 research credits in addition to a $20 Amazon gift card for participation. A signed consent form providing experiment run-time, task demands and risks, and the decision to quit at any time was collected. Three participants were rejected from the final analysis, leaving a total sample size of 30 participants. Reasons for rejection were early termination of the study (n = 1), experimenter error (n = 1), and failure to maintain fixation throughout the experiment (n =1).

The mean age of the sample was : 21.8 years (min: 18, max: 34), gender identity was 63% female, 27% male, 10% non-binary and racial/ethnic identity as 33% non-Hispanic White, 27% Latinx/Hispanic, 20% Asian, 20% mixed-race.

### Stimuli

All stimuli and the experimental tasks were coded in MATLAB (The MathWorks Inc., 2022) using the Psychophysics Toolbox 3 (Brainard, 1997). The experiment was presented on a gamma-corrected VIEWPixx EEG monitor (53.4 x 30 cm, 1920 x 1080 resolution, 120 Hz refresh rate) in a dimly lit room. Participants sat approximately 74 cm away from the monitor with their head stabilized on a chin rest.

Stimuli were presented on a uniformly gray screen (approx. 30 cd/m^2^) with a small central fixation point which was physically iso-luminant to the background. Two darker gray pedestals (approx. 15 cd/ m^2^) were presented during the entire steady condition and only during stimulus presentation in the pulsed condition (including stimulus absent trials). The pedestals occupied 6 by 6 degrees of visual angle (DVA) and were located in the upper right and left visual field, centered at 5 DVA above and 5 DVA to the right and left of the fixation (see Figure 1A). A solid black outline was drawn around the pedestals and was presented during both conditions for the entire task to equate spatial uncertainty.

We modeled our stimulus design closely after that of Yeshurun and Sabo (2012) who used the SPP paradigm to study spatial attention in the two pathways. The target stimulus in this study was a Gabor patch with a vertical grating of 1.8 cycle per DVA and a spatial SD of 0.5 DVA that was clipped at 4 DVA and centered within the pedestals (5 DVA above and 5 DVA to the right or left of fixation). The contrast level of the target was determined for each condition and participant separately using a thresholding procedure, which estimated the contrast required for a roughly 70% hit rate (see *Procedure*).

### Procedure

#### Main experiment

Participants performed 720 trials of a Yes/No detection task with confidence ratings. The experiment had two conditions, termed “steady” and “pulsed,” which were counterbalanced across participants.

Participants completed a total of 10 experimental blocks; two practice blocks (one per condition), two thresholding blocks (one per condition), and six task blocks (three per condition). Simultaneous EEG was collected during the task. In all task blocks the probability of stimulus presence and the stimulus location (left/right) were randomly chosen on each trial with a 50/50 probability.

The participant’s goal was to report whether or not they had seen the stimulus on each trial and provide a confidence rating of this judgment. Detection and confidence reports were collected through a single button press and consisted of a 4-point confidence scale for both seen and unseen judgements (1 = I’m guessing I saw/didn’t see it, 4 = I’m certain I saw/didn’t see it). Participants were instructed to keep their gaze on a central fixation point throughout the task.

*Steady condition.* The steady condition has two solid, dark gray boxes (pedestals) above fixation to the right and left (see Figure 1A) where the stimulus could be presented. The difference in contrast between the background and pedestals is believed to drive a sustained response in the PC pathway, biasing the detection of the stimulus towards the MC pathway.

*Pulsed condition.* The pulsed condition has two empty frames in the upper right and left of fixation (see Figure 1A). During the 25ms period when the stimulus could be presented, the two empty frames pulse a solid, dark gray with the stimulus drawn inside one of them on stimulus-present trials. The sudden onset of the pedestals and target are thought to saturate the MC pathway cells, biasing detection to the PC pathway.

The thresholding blocks were used to determine the difficulty (via stimulus contrast level) of the subsequent task blocks so that participants had an average detection rate of approximately 70% in both tasks. We achieved this using a 1-up/2-down staircase (Kingdom & Prins, 2010) procedure, which updated the contrast level on each trial depending on the participant’s detection responses (for stimulus-present trials). For each trial that the participant reported not seeing the target, the following trial would become easier (“up”) by increasing the contrast level of the target by a small value. For every two consecutive trials a participant reported seeing the target, the following trial became more difficult (“down”) by decreasing the contrast level by the same small value. At the end of the block, the contrast levels of the last 20 reversals were averaged to obtain an individual’s threshold value.

### Pilot study

Prior to running the main experiment study, we conducted a manipulation check (n= 9) of the SPP paradigm in order to verify that the steady and pulsed conditions displayed spatial frequency tuning compatible with known properties of the MC and PC pathways. Specifically, we varied the spatial frequency of the Gabor to either align with the feature preferences of the MC pathway (1.8 cpd) or the PC pathway (6.5 cpd), keeping all else constant. To estimate detection thresholds, we varied the contrast level of the Gabor to span the psychometric function and fit a cumulative normal distribution to the hit rate data. From the fits, we estimated 50% detection thresholds for each condition and spatial frequency.

Participants completed one block of each condition, counterbalanced across participants.

### EEG acquisition and preprocessing

Continuous EEG data was recorded in Brain Vision Recorder using a 64 Ag/AgCl gel-based active electrode system (actiCHamp Plus, Brain Products GmbH, Gilching, Germany). EEG data were processed using a custom MATLAB script (version R2022a) and the EEGLAB toolbox (Delorme & Makeig, 2004). We began by applying a high-pass filter (0.1 Hz) to the data, downsampling to 500 Hz, and re-referencing to the median of all electrodes. The data were then epoched into 4-second windows centered on stimulus onset and visually inspected for quality. The rejection criteria for data cleaning included excessive noise, muscle artifacts, or ocular movement (e.g., blinks) around -500 to 200ms pre- and post-stimulus. An average of 125 trials (17%) were rejected (min: 45, max: 198). Channels with excessive noise were removed and spherically interpolated (mean: 5.4, min: 1, max: 9). Finally, the data were average referenced.

### Pre-stimulus alpha power

Single-trial pre-stimulus alpha power was estimated for each electrode using a fast Fourier transform (FFT) on a 500 ms pre-stimulus window. The pre-stimulus signal was Hamming-tapered and zero padded (by a factor of 5) to increase frequency resolution. Power was obtained by squaring the absolute value of the FFT, after which a log10 transform was applied in order to help normalize skew in single-trial alpha power (Smulders et al., 2018). Frequencies past the Nyquist frequency were removed.

Alpha power was averaged over a cluster of eight occipital electrodes which demonstrated the highest group-level alpha-band power (see Figure 1D) as well as over an individual’s peak alpha frequency (IAF) +/- 2 Hz. IAF was identified using the MATLAB function (findpeaks.m) which assigned a given participant’s IAF as the frequency with the largest power in the trial-averaged power spectrum in the range 7-14 Hz (mean IAF = 9.9 Hz). For any participant who did not show a discriminate peak in the power spectrum in the alpha-band range, an IAF of 10 was assigned (n = 3). Once a single estimate of alpha power was assigned to each trial, the median alpha power value was assessed for each condition. This value was used to designate strong (trials whose power was above the median) and weak (below the median) alpha power trials.

For analyses conducted on power spectra, alpha power estimates were computed by taking the average alpha power of the power spectrum across the IAF +/- 2 Hz. Due to some participants not having any false alarms (n=3), the total sample size for false alarm versus correct rejection analyses is 27.

### Signal detection measurements

*Criterion.* Criterion was calculated by adding the z-transformed hit rate (HR) to the z-transformed false alarm rate (FAR) and multiplying the sum by -1/2.

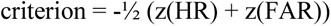

*D-prime.* D’ is calculated by subtracting the z-transformed FAR from the z-transformed HR.

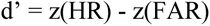

A loglinear correction was applied to the HR and FAR, which assumes a small number of false alarms or misses given an infinite number of trials (Stanislaw & Todorov, 1999).

*Generalized linear model (GLM).* To confirm the results of our median-split analysis using a single-trial approach, we additionally estimated SDT parameters in a GLM with single-trial pre-stimulus alpha power as a factor. Specifically, the coefficients from a probit-link function GLM can be used to compute SDT measurements (DeCarlo, 1998). The model estimates the probability of the responding “seen” (resp) given stimulus presence (stimpres), pre-stimulus alpha power (alphapow), and the interaction term of the two (stimpres:alphapow).

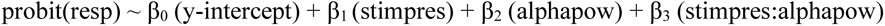

*GLM detection reports*. The first two terms are used in computing detection. For stimulus absent trials (when β_1_ equals 0) β_0_ represents the FAR. During stimulus present trials, β_0_+ β_1_ represents the HR.

*GLM criterion.* C is computed by taking the negative of the sum of the y-intercept and alpha power coefficient and subtracting that by half of the sum of the stimulus presence coefficient and the interaction coefficient.

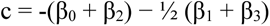

*GLM alpha power on criterion.* The effect of alpha power on criterion is computed by subtracting the negative of the alpha power coefficient from half of the interaction coefficient.

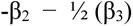

*GLM d-prime.* D’ is calculated by adding the stimulus presence coefficient to the interaction coefficient.

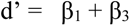

*GLM alpha power on d-prime.* The effect of alpha power on sensitivity is the interaction coefficient, β_3_.

## Results

### Pilot behavior

We examined changes in threshold detection between the steady and pulsed condition using both a low and high spatial frequency Gabor. Our hypothesis is that the threshold for detection of a low spatial frequency Gabor will be lower in the MC-mediated steady condition compared to pulsed. Furthermore, this effect should flip for the high spatial frequency Gabor, such that threshold detection will be lower for the PC-mediated pulsed condition compared to steady. By varying stimulus contrast and fitting a psychometric function to the detection reports as a function of contrast to estimate 50% detection thresholds, we indeed found a statistically significant interaction effect in the predicted direction (F(1,8) = 9.26, p = .016; see Figure 1B), suggesting that the pedestal manipulation between the two conditions is sufficient to bias detection to one pathway.

### Main experiment behavior

In the main experiment, participants underwent a thresholding block for each condition in order to match approximately 70% hit rate. Using a paired-samples t-test we confirmed that the hit rate was indeed statistically the same between the steady and pulsed condition (t(29) = -0.93, p = 0.36, 95% CI: [-0.07, 0.02]), with an average hit rate of 66% (SD: 10%; min: 45%; max: 98%) in the steady condition and 68% (SD: 11%; min: 44%; max: 88%) in the pulsed. However, the two conditions did vary in false alarm rate (t(29) = -3.13, p = .004, 95% CI: [-0.10, -0.02]), indicating that in order to obtain the same near-threshold hit rate a higher false alarm rate was needed in the pulsed condition (M: 13%; SD: 09%; min: 01%; max: 41%) compared to the steady (M: 07%; SD: 05%; min: 01%; max: 20%).

Prior research investigating the SPP paradigm indicates an expected difference in detection threshold levels between the steady and pulsed conditions, with higher thresholds needed for pulsed when using a low-spatial frequency Gabor (Leonova, et al., 2003; Pokorny, 2011). Our main experiment replicated this pattern with a significant increase in the threshold level for the pulsed condition compared to steady (t(29) = 10.11, p < 0.001, 95% CI: [0.05, 0.08]; see Figure 1C).

### Pre-stimulus alpha power on detection

The goal of this study was to better understand how spontaneous, pre-stimulus alpha power affects magno- and parvo-mediated detection. To address this question, participants performed a detection task while we measured single-trial alpha power preceding detection and confidence reports. Using paired-sample t-tests, we analyzed the power spectra differences in the alpha-band range for hit (H) versus miss (M) trials and false alarm (FA) versus correct rejection (CR) trials.

As seen in Figure 2A, we found a significant difference in pre-stimulus alpha power for H versus M trials in the steady condition (t(29) = -2.72, p = .011, 95% CI [-0.06, -0.01]), which survives a Bonferroni correction of p < 0.0125, but no significant difference for FA versus CR trials (t(26) = 0.87, p = .39, 95% CI [-0.04, 0.10]). Conversely, we did not observe a significant difference in alpha for H versus M trials in the pulsed condition (t(29) = 1.60, p = .12, 95% CI [-0.01, 0.04]), nor for FA versus CR trials (t(26) = - 0.49, p = .63, 95% CI [-0.09, 0.05]). The finding that alpha power is generally lower for H versus M trials is a well-replicated observation (for a review: Iemi et al., 2017), however our study suggests a nuance to this effect. The effects of alpha on detection for H and M trials seem to primarily be driven by detection mediated by the MC pathway. To confirm this interaction effect, we conducted a 2-by-2 repeated-measures ANOVA with response type (hit, miss) and condition (steady, pulsed) as predictors of alpha power. Indeed, this analysis showed a significant interaction effect (F(1,29) = 10.97, p = .003) and no main effect of response type (F(1,29) = 0.09, p = .81) or condition (F(1,29) = 5.78, p = .09).

**Figure 2:**
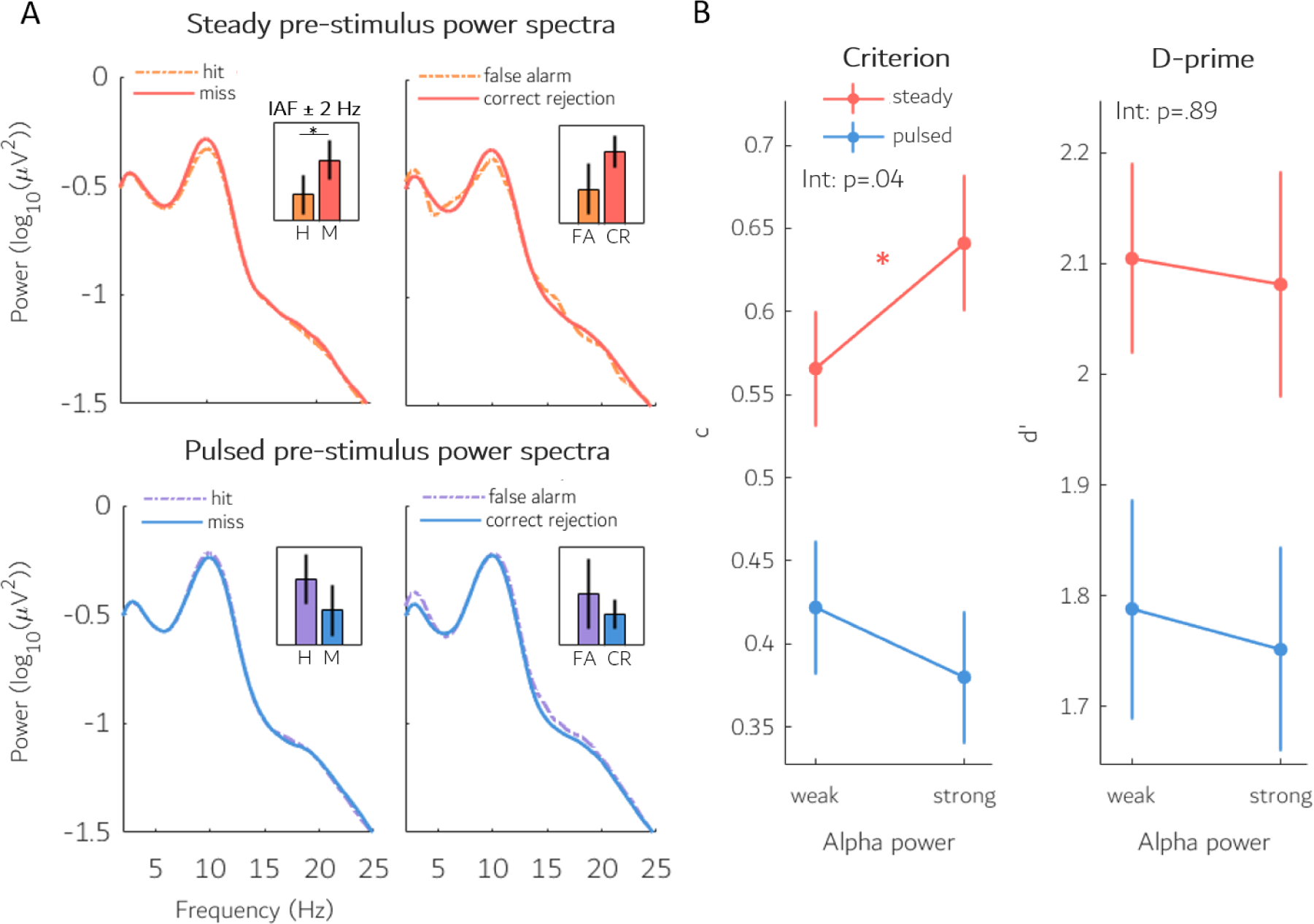
**A**) The pre-stimulus power spectra for hit (H), miss (M), false alarm (FA), and correct rejection (CR) trials for the steady and pulsed conditions. In the steady condition, there was significantly lower alpha power preceding hits versus misses, and a similar, though non-significant difference between FA and CR. In the pulsed condition, however, there were no significant effects of alpha on H versus M or FA versus CR. This suggests that the typically reported effect that alpha has on detection (namely, states of weak pre-stimulus alpha power lead to increased detection) is primarily driven by MC-mediated detection. **B**) Grand average criterion and d’ as a function of alpha power. There was a significant shift in criterion for the steady condition, but not pulsed, resulting in more conservative judgements for states of strong alpha power in the MC-mediated steady condition. We observed no significant effect of alpha power on sensitivity in either condition. Error bars are +/- 1 SEM.

### Effects of pre-stimulus alpha power on criterion, d’, and confidence

After revealing a selective effect of pre-stimulus alpha power on MC-mediated detection, we further investigated how detection was changing using the signal detection framework. This allowed us to assess whether changes in detection were arising from a shift in response bias (criterion) or a change in sensitivity (d’). We analyzed changes in criterion and d’ as a function of alpha power (weak, strong) and condition (steady, pulsed) using a 2-by-2 repeated-measures ANOVA.

As shown in Figure 2B, we found a no significant main effect of alpha power on criterion (F(1,29) = 0.43, p = .52), a significant effect of condition (F(1,29) = 13.55, p < .001) and, critically, an interaction effect of alpha with condition (F(1,29) = 4.77, p = .04). Follow-up t-tests showed a significant shift in criterion for the steady condition (t(29) = 2.20, p =.04, 95% CI: [0.01, 0.15]), but not pulsed (t(29) = -1.05, p = .30, 95% CI: [-0.12, 0.04]). This shows that the change in detection for the MC-mediated steady condition was the result of adopting a more liberal detection criterion during states of weak alpha power. It also suggests no effect on criterion for the PC-mediated pulsed condition.

There was a significant effect of condition on d’ (F(1,29) = 4.76, p = .04) but no main effect of alpha on d’ (F(1,29) = 0.41, p = .53) nor an interaction (F(1,29) = 0.02, p = .89). This suggests that alpha power does not influence perceptual sensitivity in either MC- or PC-mediated detection.

Our next goal was to assess how spontaneous states of pre-stimulus alpha power may be affecting confidence ratings. To understand how confidence ratings might change along with alpha-induced criterion shifts, we first simulated confidence responses in a SDT framework (Figure 3, left-side images). To this end, we assumed that internal responses of stimulus present and stimulus absent are each drawn from Gaussian distributions with equal variance but different means (in particular, a higher mean on stimulus present trials). Observers then compare single-trial samples of internal evidence to a decision criterion to decide if the stimulus was present or not. Because observers in our study were conservative (i.e., criterion > 0), we ran our stimulation with a conservative criterion. In this framework, confidence is simply the distance that a sample of evidence on a given trial is from the decision boundary, such that a sample of evidence far from the criterion (in either direction) would be endorsed with high confidence and a sample near the criterion would be endorsed with low confidence. To understand how alpha might change confidence, we assumed that states of low pre-stimulus power cause an additive shift in the internal response distributions (Pilipenko & Samaha, 2024), such that both the signal and noise distributions are shifted to the right, while the criterion stays in the same location. This captures the empirical finding in our study that, at least in the steady condition, observers were more liberal when pre-stimulus alpha was low, with no change in sensitivity. The simulation revealed that, if confidence is understood as distance to the criterion and subjects have a conservative criterion, then decreasing both distributions by the same factor leads to higher confidence for “unseen” trials (trials that did not surpass the criterion) but lower confidence for “seen” trials. Since we further expected this interaction effect only in the MC-mediated steady condition of our experiment, we therefore assessed changes in confidence as a function of alpha power (weak, strong), response (seen, unseen), and condition (steady, pulsed) using a 2- by-2-by-2 repeated-measures ANOVA.

**Figure 3:**
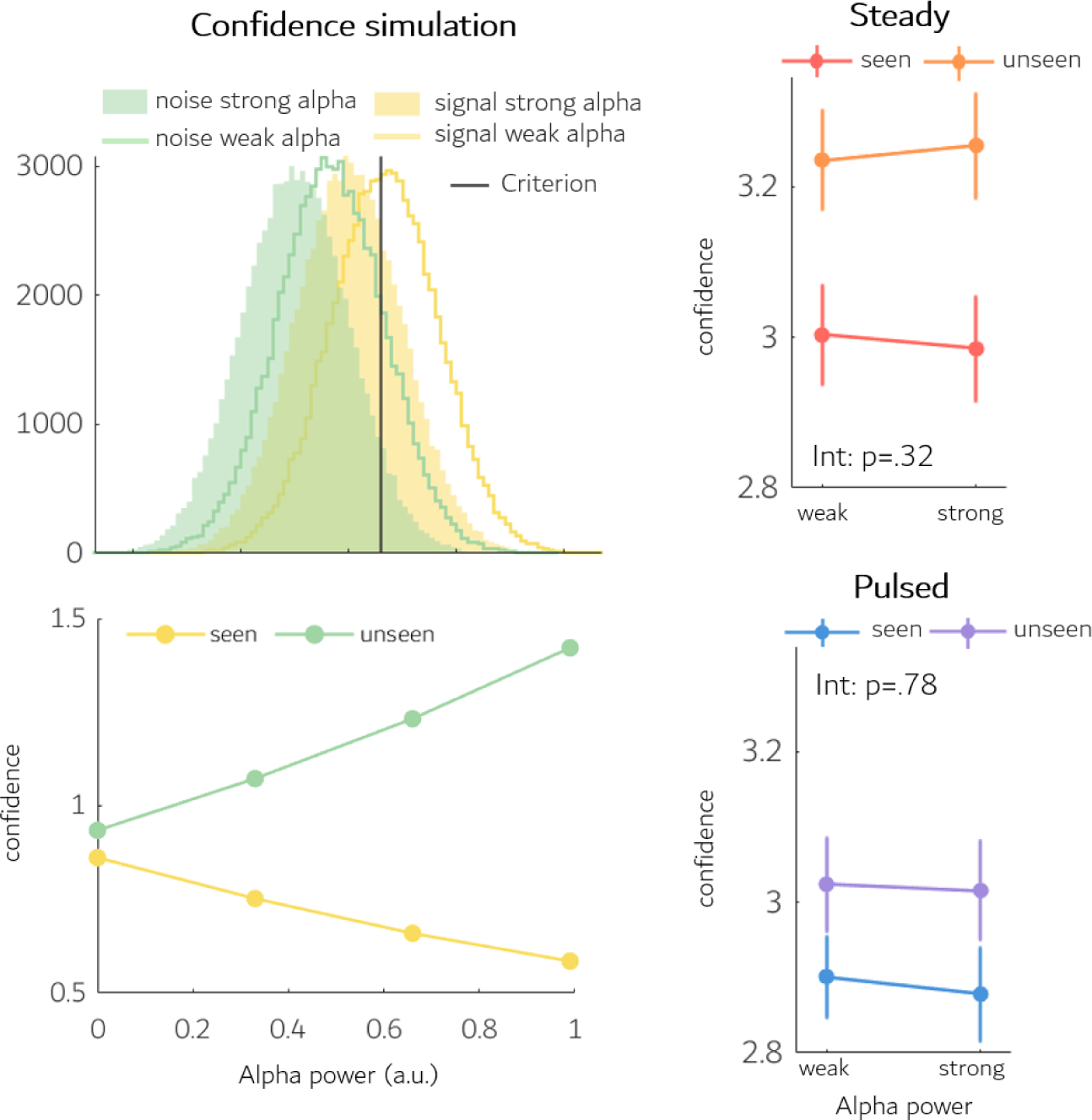
The simulation shows the distributions of signal (yellow) and noise (green) for weak (line) and strong (filled) alpha power. The additive effect of weak alpha power shifts both distributions by the same constant value, altering their position relative to the decision criterion. Confidence is calculated as a function of distance from criterion with higher confidence being farther away from criterion. As alpha power increases, the distributions shift to the left, increasing confidence for unseen trials (left of criterion) and decreasing confidence for seen trials (right of criterion). While these predictions were not significantly replicated in our confidence reports, our observed shifts in criterion may have been too modest to detect these effects in confidence. Error bars are +/- 1 SEM.

There was a significant main effect of response on confidence (F(1,239) = 9.17, p = .005), as predicted by our simulation, and a trending, though not significant, effect of condition (F(1,239) = 3.85, p = .060). This supports a significant change in confidence depending on whether or not the stimulus had been seen, which is predicted by the farther distance of the distributions on the left (unseen) side of the criterion compared to the right (seen), given a conservative criterion (see Figure 3). However, we observed no significant main effect of alpha power on confidence (F(1,239) = 0.3, p = .59), no main effect of alpha on response (F(1,239) = 0.55, p = .46), nor the predicted 3-way interaction of alpha by response by condition (F(1,239) = 0.12, p = .74). These results do not support a significant modulation of alpha power on confidence in either condition for ‘seen’ and ‘unseen’ responses. Although the pattern of mean confidence in the steady condition are in line with those predicted by the simulation, the lack of statistical significance could be due to the relatively small difference in the criterion between weak and strong alpha (steady: -0.07, pulsed: 0.04) which is expected to lead to only a modest shift in confidence ratings, and thus may be too subtle to detect in this data.

### GLM alpha power

Our primary analysis dichotomizes an otherwise continuous alpha power estimate into either a “weak” or “strong” bin. In order to consider the continuous alpha estimates, we used a generalized linear model (GLM) to compute analogous SDT measurements (DeCarlo, 1998; see *Materials and Methods*). These results reflected our findings in the primary analysis suggesting an asymmetric effect of alpha on criterion which emerges during MC-mediated detection.

This approach revealed a statistically significant difference in criterion between the steady and pulsed condition (t(29) = 2.77, p = .01, 95% CI: [0.09, 0.61]) and no difference in d’ (t(29) = 1.21, p = .24, 95% CI: [-0.23, 0.91]). When assessing how alpha power affected criterion within each condition, we found a significant effect of alpha on criterion in the MC-mediated steady condition (t(29) = 2.14, p = .04, 95% CI: [0.004, 0.177]) but not the parvo-mediated pulsed condition (t(29) = -0.69, p = .50, 95% CI: [-0.16, 0.08]). This supports the selective criterion effects in the steady condition which we observed in our primary analysis. When looking at the effects of alpha power on d’ within the conditions, we found no evidence of an effect of alpha power on d’ in either the steady (t(29) = -0.76, p = .45, 95% CI: [-0.28, 0.13]) or pulsed (t(29) = -0.25, p = .81, 95% CI: [-0.20, 0.15]) condition. Finally, when assessing the interaction that alpha power had on criterion and d’ between the two conditions, we found a trending, but nonsignificant, difference of alpha on criterion depending on the condition (t(29) = 1.79, p = .08, 95% CI: [-0.02, 0.28]) and no difference in d’ (t(29) = -0.40, p = .69, 95% CI: [-0.33, 0.23]). The results from this approach generally support our primary findings, suggesting that our binning procedure did not induce spurious results.

## Discussion

This study investigated the effects of pre-stimulus alpha power on magno- and parvo-mediated detection. To examine this, we adapted the Steady versus Pulsed Pedestal paradigm as a Yes/No detection task and recorded simultaneous EEG. Our results suggest that alpha power does not impact all visual information indiscriminately and instead has selective effects on MC-mediated detection, manifesting as a liberal shift in criterion for states of weak alpha power and no change in sensitivity.

The finding of a selective effect of ongoing alpha power on MC-mediated processing is supported across the literature. Indeed, many alpha-related studies deploy a stimulus whose features align with MC-mediated detection, including low spatial frequencies (Iemi et al., 2017; Samaha et al., 2017; Zhou et al., 2021; Pilipenko & Samaha, 2024), short presentation duration (Ergenoglu et al., 2004; Samaha et al., 2022; Benwell et al., 2022), and being peripherally located (Babiloni et al., 2006; Thut et al., 2006; Di Gregorio et al., 2022). The general findings in these studies support the inhibitory account of alpha as seen in our MC-mediated steady condition with weaker alpha power preceding hits compared to misses. Counter to this canonical magno-centric stimulus, one study conducted by Samaha et al. (2022) utilized a naturalistic stimulus set in order to understand alpha’s effects on high-level perception. Participants were asked to discriminate between images of faces and houses, a task which could be argued to recruit the ventral visual stream. The results of this study showed an increase in subjective visibility judgments during weak alpha with no accompanying change in accuracy – evidence which is taken in support of alpha’s regulation of cortical excitability, even during high-level perception. However, the stimulus set in this study did not control for low-level visual features, such as spatial frequency content, which are pathway-specific. Therefore, it is still expected that both visual pathways were involved in the task. Taken together, the literature suggests that most reported studies have not yet assessed the effects of alpha on PC-mediated detection and that pre-stimulus alpha power may be MC-pathway specific.

Inferred support for the lack of alpha effects in the PC-mediated pulsed condition can be taken from Iemi et al. (2022), who looked at the relationship between oscillatory alpha activity and broadband high-frequency activity (BHA; 70-150 Hz) across the human cortex using intracranial recordings. The significance between these two metrics is that spontaneous alpha oscillations are thought to reflect varying levels of cortical excitability (with strong alpha indexing low cortical excitability) whereas BHA is considered a proxy for neuronal spiking activity (with greater BHA indicating more neural firing).

Thus, the two are expected to covary with a negative relationship if oscillatory alpha activity is related to cortical excitability and increased firing. The anatomical map of the relationship strength shows that the correlation is largely skewed towards the dorsal stream compared to the ventral stream, which generally shows a minimal, if not no, correlation. This could indicate that the functional role of alpha may in part be pathway specific, with its relationship to cortical arousal manifesting from the MC-pathway.

One other notable study whose results complement ours looked at the effects of attention on alpha oscillations in the ventral stream. Mo et al. (2011) recorded intracranial local field potentials from the inferotemporal cortex (IT) of two macaque monkeys while they attended either to an audio or visual stimulus during an “odd one out” task. Since the IT cortex, as part of the ventral stream, is thought to have greater contributions from the PC pathway, how alpha behaves in this region could inform the functional role of alpha along more PC-biased versus MC-biased cortical areas. Indeed,while performing the visual-attention task, they found that alpha power was *higher* while attending to the visual stimuli compared to ignoring the visual stimuli. This goes against the typical findings, where there is a decrease in alpha for visually attended stimuli (Worden et al., 2000; Liu et al., 2014; Wan et al., 2019). Thus, the findings reported by Mo et al. suggests that alpha’s role in perception might be opposite in ventral stream areas, which could help explain the trending conservative change of criterion during states of weak alpha we observed in the pulsed condition.

We had expected to observe an impact of pre-stimulus alpha power on participants’ confidence reports based on the SDT model of confidence and the BSEM theory of alpha. Specifically, we assumed that states of strong alpha power produce an additive shift in both the signal and noise distribution, shifting them around the internal decision criterion. If confidence is computed by considering the distance of a given piece of evidence from the decision criterion, the shifted distributions would lead to a pattern where the average confidence ratings would increase for unseen trials and a decrease in seen trials (see Figure 3). However, we did not find evidence for this prediction despite robust findings indicating that weak alpha boosts both confidence (Samaha et al., 2017; Pilipenko & Samaha, 2024) and subjective visibility judgements (Benwell et al., 2017; Benwell et al., 2022). This lack of a measurable effect may be due to relatively modest shifts in cortical excitability within this dataset, as inferred by the size of the criterion shift between strong and weak alpha power. Increasing the number of task blocks could potentially expand the range of alpha power, helping to reveal significant effects of alpha power on confidence judgments.

The results of this study contribute to our understanding of the functional role of alpha oscillations in conscious visual perception. Understanding how alpha power differentially influences the processing of various features may help understand the diversity of functional roles supported by alpha oscillations and how these vary across cortical locations and/or distinct alpha generators. Further research should consider systematically varying low-level stimulus properties such as spatial frequency, visual field location, and color as well as use paradigms which encourage pathway-specific stimulus processing in order to replicate and expand these results.

## Acknowledgements

We sincerely thank Jessica De La Torre, Vrishab Nukala, Maxwell Volkan, Marcella Williams, Montana Wilson for help with data collection.

